# Characterization of the cytopathic effect of monkeypox virus isolated from clinical specimens and differentiation from common viral exanthems

**DOI:** 10.1101/2022.09.13.507875

**Authors:** Angela Ma, Janine Langer, Kimberly E. Hanson, Benjamin T. Bradley

**Affiliations:** Department of Pathology, University of Utah, Salt Lake City, UT; ARUP Laboratories, Salt Lake City, UT; Division of Infectious Diseases, Department of Medicine, University of Utah, Salt Lake City, UT

**Keywords:** monkeypox virus, MPXV, viral culture, cytopathic effect, viral load, virology

## Abstract

While the practice of viral culture has largely been replaced by nucleic acid amplification tests, circumstances still exist in which the availability of viral culture will allow for the diagnosis of infections not included in a provider’s differential diagnosis. Here, we examine the cytopathic effect (CPE) and clinical data associated with eighteen cases of monkeypox virus (MPXV) isolated from nineteen clinical samples submitted for viral culture. During the study period a total of 3,468 viral cultures were performed with herpes simplex virus most commonly isolated (646/3,468; 18.6%), followed by monkeypox virus (19/3,468; 0.6%) and varicella zoster virus (12/3,468; 0.4%). Most MPXV-positive samples were obtained from males (14/19) and taken from genital (7/19) or rectal lesions (5/19). Cycle threshold values of tested samples ranged from 15.3 to 29.0. Growth of MPXV in cell culture was rapid, yielding detectable CPE at a median of 2 days (range: 1-4) often with >50% of the monolayer affected in RMK, BGM, A549, and MRC-5 cell lines. As clinical features of MPXV, HSV, and VZV lesions may overlap, CPE patterns were comparted between viruses. HSV CPE developed in a similar time frame (median: 2 days, range: 1-7) but was more often negative in RMK cells relative to MPXV. VZV grew more slowly (median: 9 days, range: 5-11) and demonstrated CPE affecting ≤25% of cell monolayers when positive. Viral culture remains an important tool for the detection of rare or emerging viral pathogens, particularly when high viral load specimens are easily obtained.

## Introduction

The 2022 monkeypox virus outbreak has rapidly evolved since the first confirmed case in a United Kingdom traveling returning from Nigeria on May 7^th^, 2022 (1, 2). Following evidence of sustained transmission in non-endemic countries, the outbreak was declared a Public Health Emergency of International Concern by the World Health Organization and US Department of Health and Human Services (3, 4). Monkeypox virus (MPXV) is a member of the *Orthopoxivirus* genus, which is comprised of other human pathogens including cowpox, vaccinia, and variola viruses, the latter being the causative agent of smallpox disease (5). Human infection with MPXV was first recorded in 1970, and although the zoonotic reservoir remains unknown, rodents are suspected to be the most likely source (6). Two phylogenetic clades of MPXV are recognized, Congo Basin (Central African) and West African (7). Infection with the Congo Basin lineage is associated with more severe illness resembling smallpox and is classified as a select agent by the CDC, whereas West African monkeypox infection typically demonstrates decreased clinical severity and transmission (8). The current monkeypox outbreak has been attributed to the West African clade (1). Predominantly endemic to regions of Africa, sporadic cases of travel-associated monkeypox have been documented in other countries without sustained human-to-human transmission. Prior to the current outbreak, the last US monkeypox outbreak was reported in 2003 and was associated with infected prairie dogs exposed to rodents originating from Ghana (9). Human-to-human transmission of monkeypox was presumed to infrequently occur; however, direct contact between individuals has been a predominant driver in the spread of the 2022 outbreak (4). Preliminary data suggests decreasing population immunity due to termination of smallpox vaccine administration and novel mutations in outbreak-associated isolates have contributed towards increased human transmission (10).

Nucleic acid amplification tests (NAATs) have largely become the primary diagnostic method of viral infections due to their rapid turnaround times and increased sensitivity relative to the historical gold standard, viral culture. While conventional viral culture is more laborious and resource intensive, its utility during novel pathogen outbreaks should not be dismissed. Viral culture can detect a wide range of pathogens not covered by common NAATs and isolate viruses that may not have been part of a provider’s original differential diagnosis. (11). Furthermore, culture growth is less impacted by strain variability or genetic mutations that may contribute to false negative results by NAATs (12). Viral culture is necessary for phenotypic antiviral susceptibility testing and culture results can be useful in determining infectivity in situations of prolonged viral shedding (13, 14). Few studies have documented viral culture of MPXV with most being limited to the Congo Basin lineage (15–17). Moreover, the appearance of cytopathic effects (CPE) caused by MPXV has been poorly characterized from the standpoint of the practicing clinical microbiologist.

At Associated Regional and University Pathologists, Inc. (ARUP Laboratories), viral cultures are routinely performed on respiratory and non-respiratory sources using shell vials. Through this technique, ARUP Laboratories identified its first case of monkeypox infection from the current outbreak on July 11^th^, 2 weeks prior to the availability of an in-house orthopoxvirus molecular assay. In this study, we examine the growth characteristics and CPE of MPXV isolated from primary clinical specimens. MPXV CPE is also compared to herpes simplex virus (HSV) and varicella-zoster virus (VZV) which may cause similar lesions.

## Methods

### Data collection

Laboratory records were reviewed to identify viral culture test results from July 11^th^, 2022 to August 26^th^, 2022. Demographic information and specimen handling timestamps were recorded. For specimens with unusual CPE that were ultimately confirmed to be MPVX positive by molecular testing, the laboratory information system was queried to see if any additional microbiological testing had been ordered in the 48 hours surrounding specimen collection. For comparison of HSV and VZV CPE to MPXV CPE, 20 HSV-positive samples and 11 VZV-positive samples were evaluated. An IRB exemption was granted by the University of Utah IRB under ID#: 00158025.

### Viral Culture

Swabs were shipped in viral transport media at approximately 4°C and received at ARUP Laboratories. Upon receipt, specimens were processed for viral culture. Briefly, the tube was vortexed to release bound cells or virus, and transport media was aliquoted into a labeled holding tube where a 0.2 mL mixture of penicillin, streptomycin, and Amphotericin B was added. The tube was then centrifuged at 1500 x g for 10 minutes to clarify the sample. Shell vials containing cell lines were each inoculated with 0.2-0.3 mL of the treated supernatant. Depending on the test ordered, individual shell vials containing rhesus monkey kidney cells (RMK) (Quidel, San Diego, CA), Buffalo green monkey kidney cells (BGM) (ARUP Reagent Laboratory, Salt Lake City, UT), A549 human lung carcinoma (ARUP Reagent Laboratory, Salt Lake City, UT), or MRC-5 human embryonic lung fibroblast cell lines (ARUP Reagent Laboratory, Salt Lake City, UT) were inoculated. Cell lines used for each assay were as follows – Non-respiratory broad viral culture with or without CMV DFA: RMK, BGM, A549, and MRC-5 cell lines; Herpes simplex viral culture: A549 cell line; Varicella-zoster viral culture: A549 and MRC-5 cell lines. Inoculated cell lines were incubated at 37°C, 5% CO_2_ in 1mL of 2% MEM Maintenance Media (ARUP Reagent Laboratory, Salt Lake City, UT). Cultures were read daily for a total of 14 days. When present, cytopathic effect (CPE) was graded as follows: 0 = no CPE present, 1+ = ≤25% CPE, 2+ = >25% CPE, 3+ = >50% CPE, 4+ = >75% CPE. Images of CPE were acquired using an inverted light microscope.

For cultures with typical CPE, diagnosis was confirmed by direct immunofluorescent (DFA) or indirect immunofluorescent (IFA) staining of the shell vial. DFA kits were used to test for herpes simplex virus types 1 and 2 (D^3^ DFA HSV Identification and Typing kit, Quidel, San Diego, CA), varicella-zoster virus (Light Diagnostics, Varicella-Zoster Virus DFA kit, EMD Millipore, Temecula, CA), adenovirus (D^3^Ultra Respiratory Virus Screening & Identification kit, Quidel, San Diego, CA), and cytomegalovirus (D^3^ DFA Cytomegalovirus Immediate Early Antigen Identification kit, Quidel, San Diego, CA). IFA kits were used to test for enterovirus (D^3^ IFA Enterovirus Identification and Typing kit, Quidel, San Diego, CA). Manufacturer instructions were followed for all kits.

Workup of cultures with atypical CPE changed over the course of the study. Initially these specimens were tested by DFA and IFA reagents to common viruses including HSV, VZV, adenovirus, and enterovirus. If workup was negative for these viruses, a presumptive diagnosis of MPXV was made. The medical directors then contacted the client to discuss ancillary molecular testing through state and local public health labs. Once laboratory staff became more familiar with the prototypical CPE of MPXV and following the availability of on-site molecular testing, cultures demonstrating CPE consistent with MPXV had their work-up stopped and were sent to orthopoxvirus NAAT for confirmation.

### ARUP Orthopoxvirus real-time PCR assay

Presumptive detection of MPXV from clinical samples was performed by a laboratory-developed real-time PCR assay using primers and probes designed by ELITech (Bothell, WA, USA) targeting a conserved region within the polymerase gene common among the *Orthopoxvirus* genus. Samples received on dry swabs were submerged in 500uL of PBS, vortexed for 15 seconds, then allowed to sit at room temperature for 1 hour before proceeding to extraction. For specimens in VTM and dry swabs following resuspension, 200uL of sample was eluted into 80ul on the Chemagic MSMI (PerkinElmer, Waltham, MA, USA) instrument. Following reaction set-up, amplification was performed on the QuantStudio 12K Flex (ThermoFisher, Waltham, MA, USA) to determine the cycle threshold. Prism (version 9.4.1, San Diego, California USA) was used for graphical and statistical analysis. An unpaired t test was performed to compare the Ct values between positive specimens sent for viral culture to those sent for orthopoxvirus NAAT only.

## Results

### Epidemiological and laboratory data for positive monkeypox virus specimens

Over the course of the study, a total of 1,482 non-respiratory broad viral cultures, 1,674 herpes simplex viral cultures, 173 varicella-zoster viral cultures, and 139 non-respiratory broad viral cultures with CMV DFA were performed. The highest positivity rate of MPXV detection was from the non-respiratory broad viral culture (14/1482; 0.9%), followed by non-respiratory broad viral culture with CMV DFA (1/139, 0.7%), varicella-zoster viral culture (1/173, 0.6%), then herpes simplex viral culture (3/1674, 0.2%). For the non-respiratory broad viral culture assay, MPXV was detected more frequently (n=14) than all other viruses (VZV, n=11; Enterovirus, n=3; Adenovirus, n=2) excluding HSV (n=170) (Table 1).

**Table 1.**
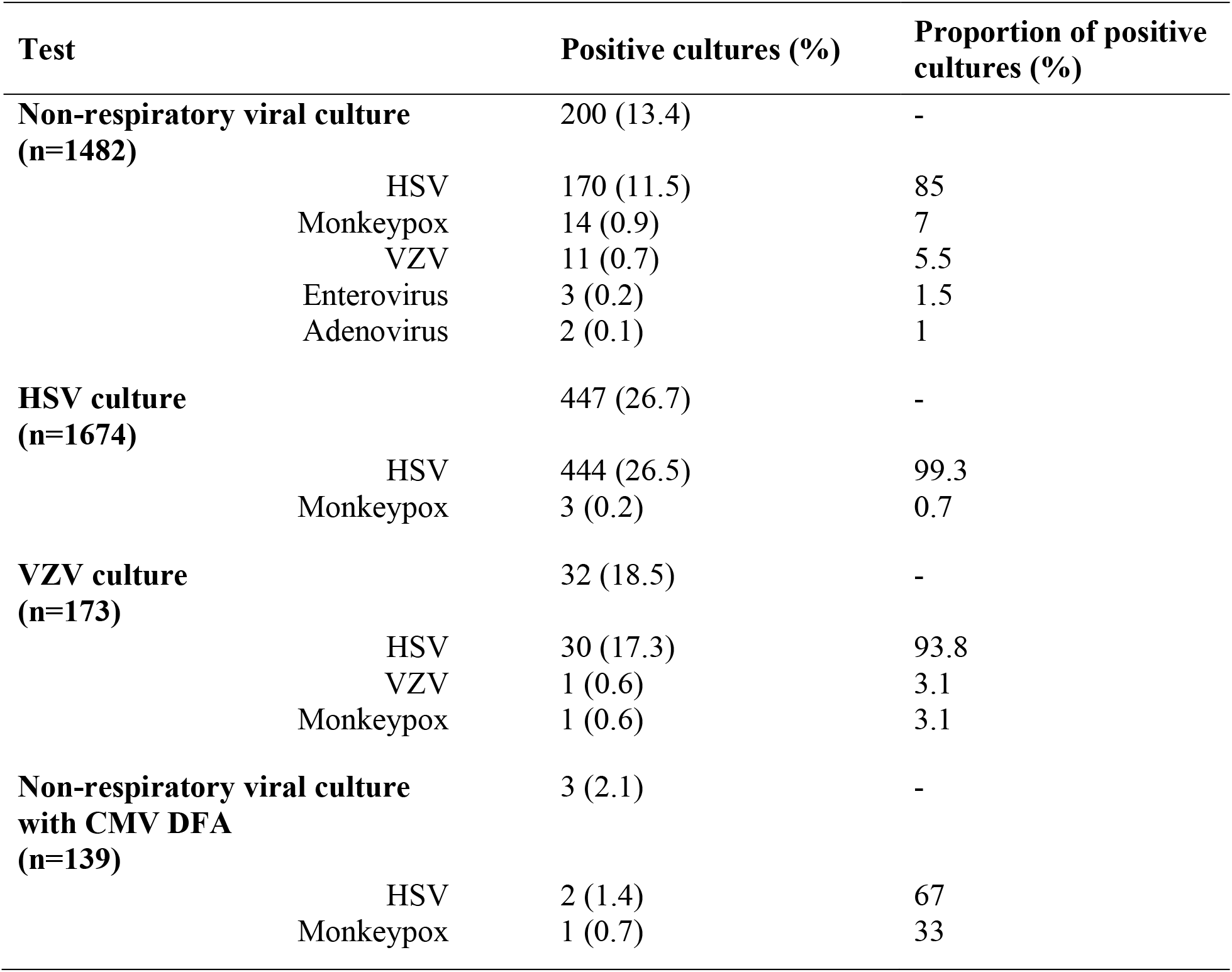
Positivity rates of non-respiratory viral culture tests performed during study period.

The 19 positive MPXV samples originated from 18 individuals including 14 males and 4 females. The median age for individuals with cell culture results suggestive of MPXV infection was 33 (range: 9 to 51 years). The predominate specimen sources were genital (7/19) and rectal (5/19). Less frequent specimen sources included skin (2/19), upper extremity (2/19), foot (1/19), pharynx (1/19), and one specimen for which a source was not provided. For many individuals (9/18), no additional infectious disease tests were requested from our lab within 48 hours of viral culture specimen collection. When additional orders were placed, they included assays to detect HSV (n=8), orthopoxvirus (n=4), and VZV (n=2). On rare occasion, testing was performed for tick-borne diseases (Table 2).

**Table 2.** Descriptive characteristics of monkeypox virus isolated from viral culture (n=19).

|  |  |  |
| --- | --- | --- |
| **Median age in years (range)** |  | 33 (9-15) |
| **Gender (n, %)** |  |  |
|  | Male | 14 (73.7) |
|  | Female | 4 (26.3) |
| **Median turnaround time in hours (range)** |  |  |
|  | Collection to receipt | 40.2 (5.2-75.4) |
|  | Receipt to result | 87.4 (29.3-214.6) |
|  | Overall time | 127.6 (53.5-256.0) |
| **Day of incubation CPE first detected (n, %)** |  |  |
|  | Day 1 | 1 (5.3) |
|  | Day 2 | 9 (47.4) |
|  | Day 3 | 5 (26.3) |
|  | Day 4 | 4 (21.0) |
| **Specimen source (n, %)** |  |  |
|  | Genital (including vagina, groin) | 7 (36.8) |
|  | Rectal (including anal) | 5 (26.3) |
|  | Skin | 2 (10.5) |
|  | Upper extremity (including hands, fingers) | 2 (10.5) |
|  | Foot | 1 (5.3) |
|  | Pharynx | 1 (5.3) |
|  | Not specified | 1 (5.3) |
| **Additional infectious disease tests ordered (n)** |  |  |
|  | No additional testing | 9 |
|  | Orthopoxvirus NAAT | 4 |
|  | HSV (including culture, DFA*, NAAT) | 8 |
|  | VZV (including DFA*, NAAT) | 2 |
|  | HIV (including 1/2 differentiation, NAAT) | 2 |
|  | AFB tissue culture | 2 |
|  | *Ehrlichia* and *Anaplasma* NAAT | 1 |
|  | *Rickettsia rickettsii* serology | 1 |
|  | Acute hepatitis serology | 1 |
| **Orthopoxvirus NAAT testing** |  |  |
|  | Cases confirmed (n, %) | 19 (100) |
|  | Average Ct value<sup>†</sup> (range) | 20.8 (15.3-29.0) |

\*Direct fluorescent antibody stain for HSV-1/2 and VZV performed on primary specimen.
<sup>†</sup>In-house testing performed on 18/19 specimens.

### Monkeypox virus is rapidly detected from viral culture

Cytopathic effect consistent with MPXV was rapidly detected from cell culture. Overall, CPE was demonstrated at a median of 2 days following inoculation. Of the 19 samples, 1 was positive at Day 1, 9 at Day 2, 5 at Day 3, and 4 at Day 4 (Table 1). Each of the four cell lines used in our assays appeared permissible to MPXV replication. Only one specimen (MPXV_0012) did not demonstrate CPE in all cell lines on the first day of positivity(Supplemental Table 1). For this specimen, both the MRC-5 and BGM cell lines were negative, but turned positive two days later. The degree of CPE varied between different cell lines with half (7/14) of the RMK cells demonstrating a CPE of 4+ while only 35% (5/14) of BGM cells had a CPE of 4+ on the first positive day (Figure 1A).

**Figure 1.**
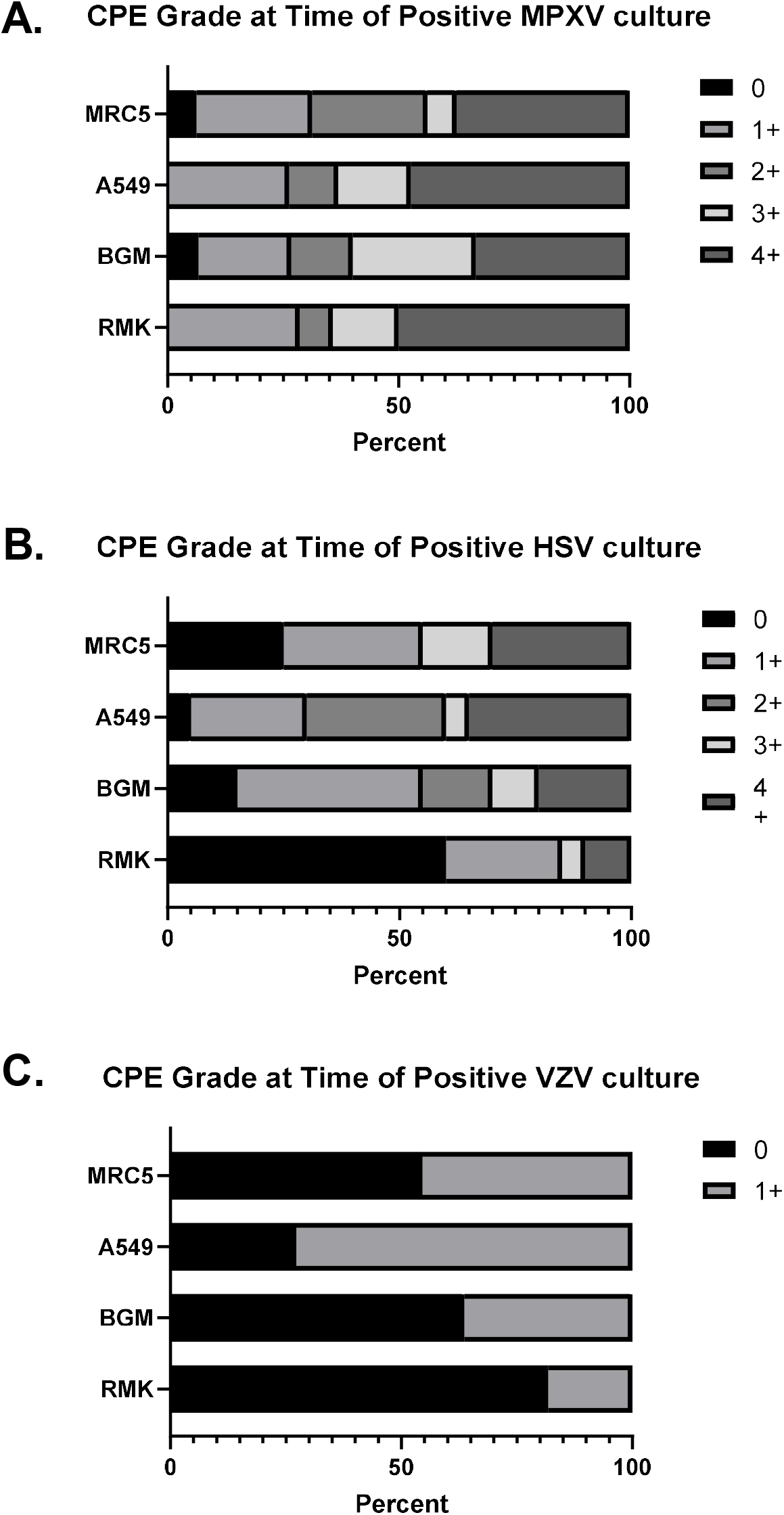
CPE Grade at Time of Positive Culture for MPXV (A), HSV (B), and VZV (C). CPE graded as follows: 0 = no CPE present, 1+ = ≤25% CPE, 2+ = >25% CPE, 3+ = >50% CPE, 4+ = >75% CPE.

Infection with MPXV led to significant, observable changes to cell culture monolayers. Uninoculated RMK cells demonstrate a biphasic population of fibroblasts and epithelial cells. When MPXV CPE was present there was a significant degree of cellular detachment and degeneration. The remaining cells were refractile and demonstrated pronounced cytoplasmic bridging. Uninoculated heteroploid BGM cells typically demonstrate a flat, cuboidal morphology, and when infected with MPXV these cells demonstrated clumping with pinching of the cytoplasmic membrane. In the A549 cells which are typically described as “cobblestone” in appearance, MPXV CPE manifested as foci of cellular detachment with surrounding cells demonstrating a glassy or “melted” appearance of their membranes. The diploid fibroblast cell line, MRC-5, grows as sheets of streaming, spindled cells in a woven appearance. MPXV CPE in these cells demonstrated cell degeneration with chords of rounded fibroblasts alternating with areas of preserved cell growth. The low-magnification appearance of streaming CPE tracks alternating with uninvolved cells was described as a “hurricane” pattern by the lab to evoke the image of the radial bands of hurricane clouds when viewed from above (Figure 2).

**Figure 2.**
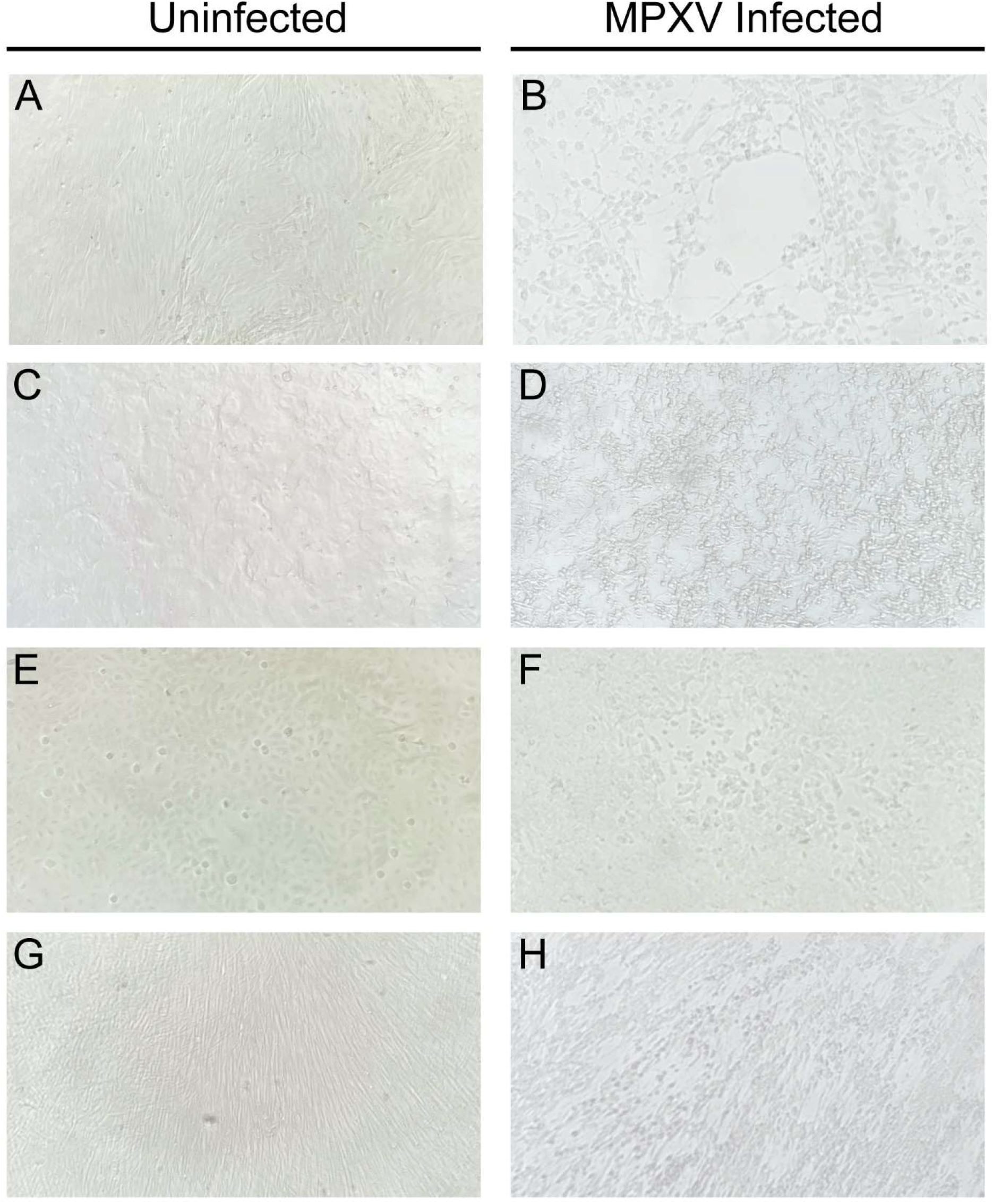
Comparison of uninfected cells with the cytopathic effect of MPXV following 2 days incubation at 37°C, 5% CO_2_ in RMK (A, B), BGM (D, D), A549 (E, F), and MRC-5 (G, H) cell lines. Images taken at 100x magnification.

### Growth characteristics and CPE patterns allow for separation of MPXV from HSV and VZV

As HSV and VZV are normally the predominate viruses isolated by culture from non-respiratory sources, the CPE of these viruses were compared to that of MPXV. A random subset of 20 HSV cases and all 11 VZV cases isolated from the non-respiratory broad viral culture assay were used for analysis. In comparison to MPXV, HSV demonstrated a similar median time to positivity of 2 days (range 1-7 days), however, a higher majority of HSV cases were positive on Day 1 in comparison to MPXV. When observable CPE was detected from HSV culture it was most frequently seen in A549 cells (19/20, 95%) followed by BGM cells (17/20, 85%). The CPE grade was lower in comparison to MPXV in all cell lines. Notably over half of RMK cells were graded as 0 for HSV while 50% (7/14) were graded as 4+ for MPXV (Figure 1A, B). VZV cultures were positive at a significantly later time, giving positive results at a median of 9 days (range 5-11 days) (Supplemental Table 1). VZV isolates demonstrated less CPE in comparison to MPXV and HSV (Figure 1C). CPE was most commonly observed in A549 cells (8/11, 73%). On the initial positive CPE read for all VZV isolates, no specimen had a CPE grade greater than 1+. As with HSV, the RMK cell line demonstrated the least amount of cytopathic effect (2/11, 18%).

In addition to growth characteristics, observable differences in the cytopathic effect of MPXV, HSV, and VZV exist. The A549 cell line potentially provides the greatest resolution as different features are observed for HSV1, HSV2, VZV, and MPXV. HSV1 and HSV2 typically present as rounded, ballooned cells with HSV2 demonstrating a wider variation in cell size (Figure 3B, C). VZV CPE in A549 cells presents as foci of less refractile, rounded cells sometimes surrounding a central clearing (Figure 3D). MPXV CPE of A549 cells also shows a central clearing though the cells have less distinct borders and a more refractile appearance (Figure 3A). For the MRC-5 cell line, the CPE pattern is similar between HSV and VZV where it presents as single or small clusters of rounded cells and contrasts to the broader patches formed by MPXV (Figure 3E, F, G, H). Within BGM cells, HSV CPE manifests as large, web-like foci of bridging cells while VZV CPE occurs in tighter foci. Similar to HSV, the CPE of MPXV occurs in patches though the foci may be more elongated. Strong CPE is infrequently observed in RMK cells infected with HSV, and in this study 10% (2/20) were observed at 4+. When present, HSV CPE in RMK cells demonstrates scattered rounded cells with refractile bodies similar to what is seen in the A549 cell line. VZV CPE in RMK presents with a central clearing surrounding by cells with cytoplasmic bridging. The CPE of MPXV in RMK cells is much more pronounced with a significantly higher rate of 4+ CPE. These cells demonstrate conspicuous cytoplasmic bridging and rapid monolayer degeneration with cells floating in the media. Descriptive findings for each virus and cell line combination are summarized in Table 3.

**Table 3.**
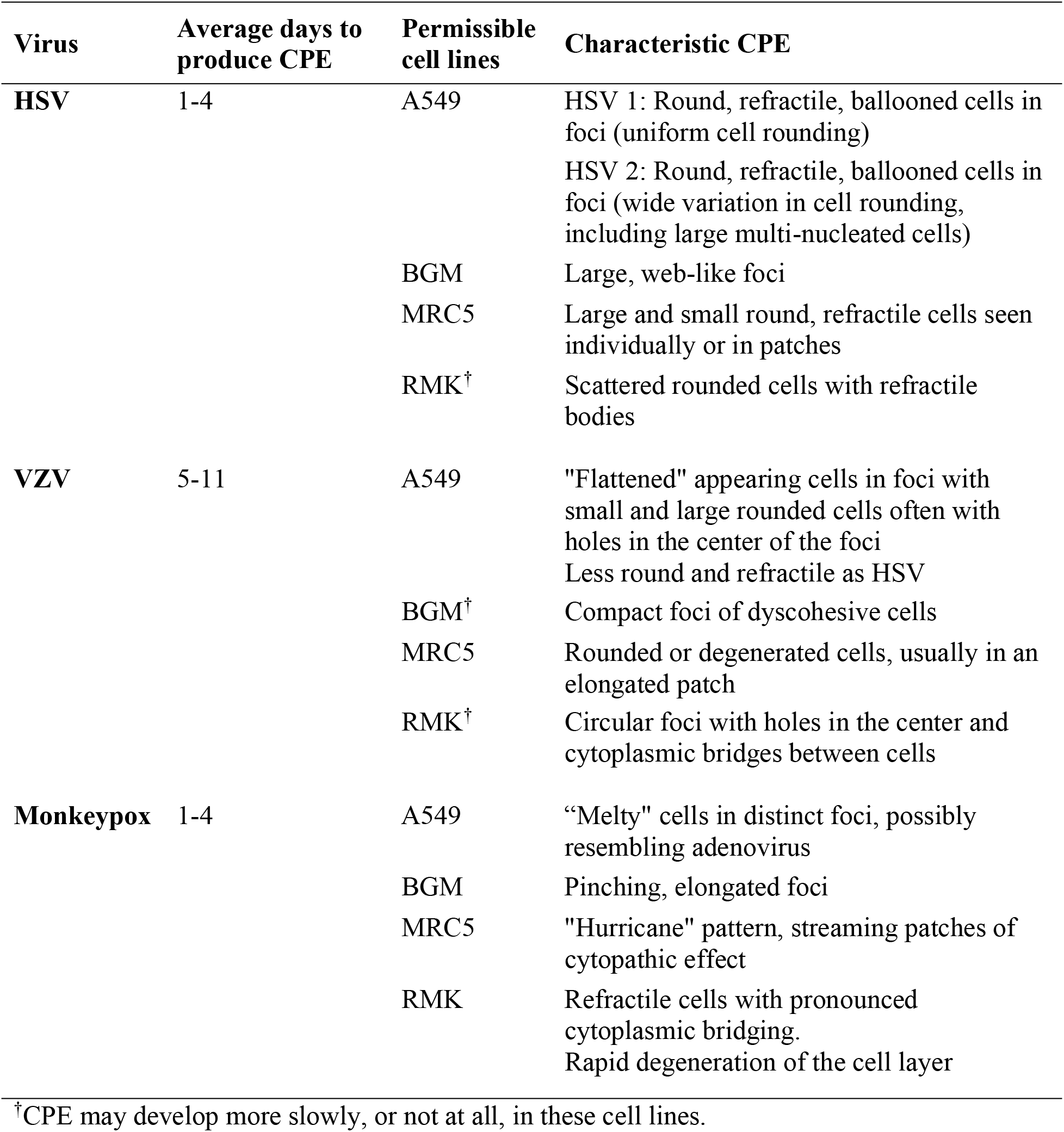
Comparison of MPXV CPE to viruses commonly isolated from non-respiratory sites.

| **Virus** | **Average days to produce CPE** | **Permissible cell lines** | **Characteristic CPE** |
| --- | --- | --- | --- |
| **HSV** | 1-4 | A549 | HSV 1: Round, refractile, ballooned cells in foci (uniform cell rounding)<br>HSV 2: Round, refractile, ballooned cells in foci (wide variation in cell rounding, including large multi-nucleated cells) |
|  |  | BGM | Large, web-like foci |
|  |  | MRC5 | Large and small round, refractile cells seen individually or in patches |
|  |  | RMK<sup>†</sup> | Scattered rounded cells with refractile bodies |
| **VZV** | 5-11 | A549 | "Flattened" appearing cells in foci with small and large rounded cells often with holes in the center of the foci<br>Less round and refractile as HSV |
|  |  | BGM<sup>†</sup> | Compact foci of dyscohesive cells |
|  |  | MRC5 | Rounded or degenerated cells, usually in an elongated patch |
|  |  | RMK<sup>†</sup> | Circular foci with holes in the center and cytoplasmic bridges between cells |
| **Monkeypox** | 1-4 | A549 | "Melty" cells in distinct foci, possibly resembling adenovirus |
|  |  | BGM | Pinching, elongated foci |
|  |  | MRC5 | "Hurricane" pattern, streaming patches of cytopathic effect |
|  |  | RMK | Refractile cells with pronounced cytoplasmic bridging.<br>Rapid degeneration of the cell layer |

<sup>†</sup>CPE may develop more slowly, or not at all, in these cell lines.

**Figure 3.**
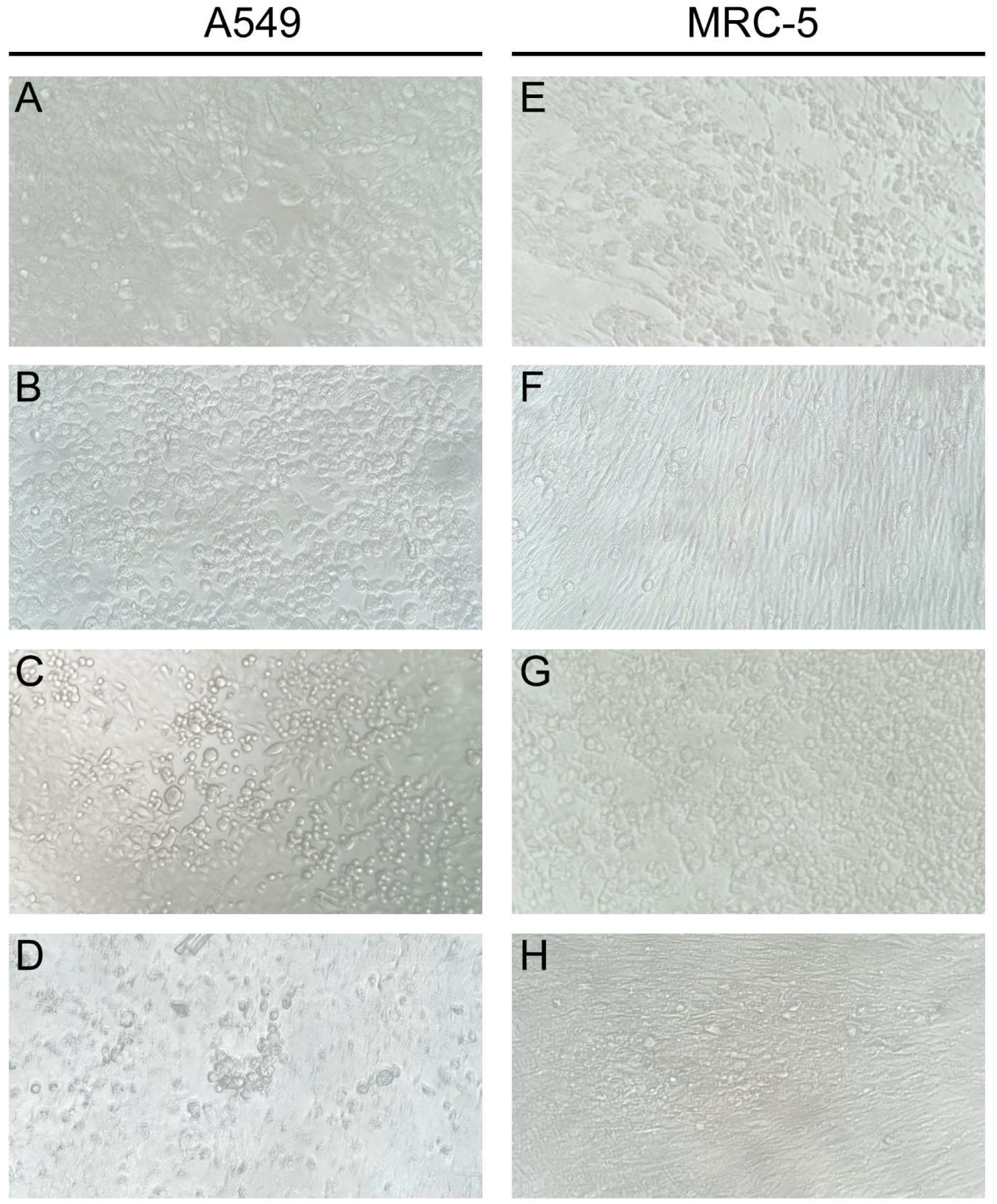
Comparison of typical MPXV (A, E), HSV1 (B, F), HSV2 (C, G), and VZV (D, H) cytopathic effect in A549 and MRC-5 cells. Images taken at 200x magnification.

## Discussion

Given the relative absence of literature, this study provides pertinent information on the CPE and growth characteristics of MPXV in common cell lines used in a clinical laboratory. We document important morphological characteristics of MPXV including rapid growth in all four cells lines, unique CPE patterns within A549 and MRC-5 cells, and the presence of strong CPE in RMK cells. These features may help laboratories that perform viral culture flag specimens as suspicious for MPXV and reduce exposure of lab personnel to high-titer virus.

The available patient characteristics in our cohort are mostly consistent with larger epidemiological trends, showing men between the ages of 20 and 50 being most affected (1). However, several outliers including a 9-year-old and 51-year-old female were also identified. In our cohort, 22% (4/18) of MPXV-positive individuals were women. This is significantly more than the 1.5% of cases reported by the Centers for Disease Control and Prevention (CDC) as of August 30^th^, 2022 (18). Perhaps due to a lower pre-test probability, the availability of viral culture may have allowed for detection of a pathogen that was not originally included in the provider’s differential diagnosis of female patients. As a reference laboratory, it is difficult to determine which other assays were included with viral culture as part of an individual patients’ work-up. However, when additional infectious disease testing was ordered at ARUP Laboratories, the differential included other viral and bacterial diseases that present with rash or ulcers (e.g., HSV, VZV, *Erhlichia*, and *Rickettsia rickettsii*).

Descriptions of MPXV CPE have mostly been limited to individual cases or passaged virus (19, 20). Whereas previous studies have shown MPXV growth in VeroE6, MA-104, LLCMK-2, OMK, and BSC-40, our study documents the CPE of direct-from-specimen MPXV in RMK, A549, BGM, and MRC-5 cell lines (15, 16, 19). Detection of MPXV CPE was rarely subtle and in the great majority of cases all cell lines demonstrated either 3+ or 4+ CPE on the day of initial positivity. While the CPE was most often strongest in RMK cells, all cell lines were capable of CPE formation. There was only one specimen (MPXV_0012) in which no CPE was observed in the BGM or MRC5 cell lines on the initial day of positivity (Supplemental Table 1). This pattern of high CPE and broad cell tropism differentiates MPXV from HSV (strong CPE but frequently limited to A549 and MRC-5 cells) and VZV (CPE slow to develop and restricted to A549 and MRC-5 cells). The elevated CPE of MPXV relative to HSV may reflect differences in viral replication kinetics or viral load; however, it may also be influenced by our limited experience in identifying MPXV from cell culture, potentially missing earlier, more subtle changes.

Evaluation of cell culture for viral CPE can be challenging and some degree of subjectivity is inherent in the method. While not all MPXV culture isolates demonstrated stereotypical CPE, we were able to identify common patterns of CPE and rapid growth kinetics. On occasion, atypical CPE would occur. In several instances, rapid cytolysis of the RMK cells prevented the identification of CPE foci. In one suspected case, rapid 3+ to 4+ CPE appeared in all cell lines but the MRC-5 cell line demonstrated fewer patches of CPE and more individually rounded cells. While suspicious, this was not definitive for MPXV. Subsequent consultation with the ordering provider revealed the patient had a disseminated HSV infection. This diagnosis was confirmed by direct fluorescent antibody testing of the shell vial for HSV.

For 18/19 specimens, the diagnosis of monkeypox virus was confirmed in our laboratory using a orthopoxvirus PCR assay. In the one case which was not tested prior to disposal, the ordering provider confirmed that the specimen tested positive for orthopoxvirus at an outside facility. Compared to specimens that were received for PCR testing only, the Ct values of viral culture samples were significantly earlier (20.8 vs. 23.9, p<0.05) highlighting the concern that culture may only detect those samples with very high viral loads. While the presence of culturable virus is not sufficient to prove infectivity, correlation between Ct values and cultivatable virus may provide a useful surrogate marker for infectiousness in epidemiological studies (21). Previous studies have performed culture when Ct values were <32 (16). Within our cohort the latest Ct observed was 29.0; however, our study was not designed to detect what the Ct limit is for cultivatable MPXV. From a diagnostic standpoint, the low Ct values observed in typical MPX lesions are reassuring that if future outbreaks were to occur, viral culture would detect a significant number of cases with the benefit of being more robust to viral mutations that may negatively affect molecular methods as has been observed with TNF receptor gene deletions (22).

Limitations of cell culture as compared to molecular techniques are its reduced ability to detect co-infections and biosafety concerns. As viral culture is performed in liquid media, individual colonies cannot be separated as is done with bacteria on solid media. In practice, this means rapidly growing viruses with wide tropism will outcompete slower, more fastidious viruses. This is of particular concern in MPXV infections where longitudinal studies in the Democratic Republic of Congo have found a 12-13% coinfection rate of MPXV with VZV in individuals presenting with monkeypox-like illnesses (16, 17). As VZV grows significantly slower than MPXV, these co-infections would likely go undetected by culture. Whether the coinfection rate is similar in the current global outbreak remains to be examined. Finally, performing viral culture places laboratory staff at risk of exposure to high-titer viral specimens. In our lab, employees were offered the JYNNEOS vaccine and specimens were handled using BSL-3 techniques in a BSL-2 environment. Because CPE is insufficient to separate the West African MPXV clade from the Congo Basin MPXV clade, which is considered a select agent under CDC guidelines, shell vials with CPE concerning for monkeypox were immediately destroyed (23). These challenges are less prevalent in molecular testing where specimens can be inactivated prior to handling.

While viral culture is no longer routinely practiced in most clinical microbiology laboratories, the unique advantages it provides help justify its continued existence within large academic medical centers and reference laboratories. Viral culture provides a more hypothesis-free method of detection for rare or emerging viral pathogens, particularly those present at high viral loads in clinical samples. In our laboratory we detected the first specimen positive for monkeypox virus on July 11^th^, 2022; two weeks prior to the availability of a molecular test in our laboratory. In seven instances where no orthopoxvirus NAAT was originally ordered, viral culture provided the first clue as to the patient’s diagnosis. Our laboratorians recognized the atypical cell CPE and alerted the medical director who then encouraged the ordering provider to add on orthopoxvirus NAAT. Thus, culture provided important diagnostic information and helped direct patient management.

## References

1. Thornhill JP, Barkati S, Walmsley S, Rockstroh J, Antinori A, Harrison LB, Palich R, Nori A, Reeves I, Habibi MS, Apea V, Boesecke C, Vandekerckhove L, Yakubovsky M, Sendagorta E, Blanco JL, Florence E, Moschese D, Maltez FM, Goorhuis A, Pourcher V, Migaud P, Noe S, Pintado C, Maggi F, Hansen A-BE, Hoffmann C, Lezama JI, Mussini C, Cattelan A, Makofane K, Tan D, Nozza S, Nemeth J, Klein MB, Orkin CM, SHARE-net Clinical Group. 2022. Monkeypox Virus Infection in Humans across 16 Countries - April-June 2022. N Engl J Med 387:679–691.

2. Minhaj FS. 2022. Monkeypox Outbreak — Nine States, May 2022. MMWR Morb Mortal Wkly Rep 71.

3. WHO Director-General’s statement at the press conference following IHR Emergency Committee regarding the multi-country outbreak of monkeypox - 23 July 2022. https://www.who.int/director-general/speeches/detail/who-director-general-s-statement-on-the-press-conference-following-IHR-emergency-committee-regarding-the-multi--country-outbreak-of-monkeypox--23-july-2022. Retrieved 22 August 2022.

4. Philpott D, Hughes CM, Alroy KA, Kerins JL, Pavlick J, Asbel L, Crawley A, Newman AP, Spencer H, Feldpausch A, Cogswell K, Davis KR, Chen J, Henderson T, Murphy K, Barnes M, Hopkins B, Fill M-MA, Mangla AT, Perella D, Barnes A, Hughes S, Griffith J, Berns AL, Milroy L, Blake H, Sievers MM, Marzan-Rodriguez M, Tori M, Black SR, Kopping E, Ruberto I, Maxted A, Sharma A, Tarter K, Jones SA, White B, Chatelain R, Russo M, Gillani S, Bornstein E, White SL, Johnson SA, Ortega E, Saathoff-Huber L, Syed A, Wills A, Anderson BJ, Oster AM, Christie A, McQuiston J, McCollum AM, Rao AK, Negrón ME, CDC Multinational Monkeypox Response Team. 2022. Epidemiologic and Clinical Characteristics of Monkeypox Cases - United States, May 17-July 22, 2022. MMWR Morb Mortal Wkly Rep 71:1018–1022.

5. McCollum AM, Damon IK. 2014. Human monkeypox. Clin Infect Dis Off Publ Infect Dis Soc Am 58:260–267.

6. Reynolds MG, Carroll DS, Karem KL. 2012. Factors affecting the likelihood of monkeypox’s emergence and spread in the post-smallpox era. Curr Opin Virol 2:335–343.

7. Likos AM, Sammons SA, Olson VA, Frace AM, Li Y, Olsen-Rasmussen M, Davidson W, Galloway R, Khristova ML, Reynolds MG, Zhao H, Carroll DS, Curns A, Formenty P, Esposito JJ, Regnery RL, Damon IK. 2005. A tale of two clades: monkeypox viruses. J Gen Virol 86:2661–2672.

8. Parker S, Nuara A, Buller RML, Schultz DA. 2007. Human monkeypox: an emerging zoonotic disease. Future Microbiol 2:17–34.

9. Centers for Disease Control and Prevention (CDC). 2003. Update: multistate outbreak of monkeypox--Illinois, Indiana, Kansas, Missouri, Ohio, and Wisconsin, 2003. MMWR Morb Mortal Wkly Rep 52:642–646.

10. Kumar N, Acharya A, Gendelman HE, Byrareddy SN. 2022. The 2022 outbreak and the pathobiology of the monkeypox virus. J Autoimmun 131:102855.

11. Martin RM, Burke K, Verma D, Xie H, Langer J, Schlaberg R, Swaminathan S, Hanson KE. 2020. Contact Transmission of Vaccinia to an Infant Diagnosed by Viral Culture and Metagenomic Sequencing. Open Forum Infect Dis 7:ofaa111.

12. Ogilvie M. 2001. Molecular techniques should not now replace cell culture in diagnostic virology laboratories. Rev Med Virol 11:351–354.

13. AlGhounaim M, Xiao Y, Caya C, Papenburg J. 2017. Diagnostic yield and clinical impact of routine cell culture for respiratory viruses among children with a negative multiplex RT-PCR result. J Clin Virol Off Publ Pan Am Soc Clin Virol 94:107–109.

14. Mileto D, Foschi A, Mancon A, Merli S, Staurenghi F, Pezzati L, Rizzo A, Conti F, Romeri F, Bernacchia D, Meroni R, Rizzardini G, Gismondo MR, Micheli V. 2021. A case of extremely prolonged viral shedding: Could cell cultures be a diagnostic tool to drive COVID-19 patient discharge? Int J Infect Dis IJID Off Publ Int Soc Infect Dis 104:631–633.

15. Hutin YJ, Williams RJ, Malfait P, Pebody R, Loparev VN, Ropp SL, Rodriguez M, Knight JC, Tshioko FK, Khan AS, Szczeniowski MV, Esposito JJ. 2001. Outbreak of human monkeypox, Democratic Republic of Congo, 1996 to 1997. Emerg Infect Dis 7:434–438.

16. Hughes CM, Liu L, Davidson WB, Radford KW, Wilkins K, Monroe B, Metcalfe MG, Likafi T, Lushima RS, Kabamba J, Nguete B, Malekani J, Pukuta E, Karhemere S, Muyembe Tamfum J-J, Okitolonda Wemakoy E, Reynolds MG, Schmid DS, McCollum AM. 2020. A Tale of Two Viruses: Coinfections of Monkeypox and Varicella Zoster Virus in the Democratic Republic of Congo. Am J Trop Med Hyg 104:604–611.

17. Hoff NA, Morier DS, Kisalu NK, Johnston SC, Doshi RH, Hensley LE, Okitolonda-Wemakoy E, Muyembe-Tamfum JJ, Lloyd-Smith JO, Rimoin AW. 2017. Varicella Coinfection in Patients with Active Monkeypox in the Democratic Republic of the Congo. EcoHealth 14:564–574.

18. CDC. 2022. Monkeypox in the U.S. Cent Dis Control Prev. https://www.cdc.gov/poxvirus/monkeypox/response/2022/index.html. Retrieved 5 September 2022.

19. Noe S, Zange S, Seilmaier M, Antwerpen MH, Fenzl T, Schneider J, Spinner CD, Bugert JJ, Wendtner C-M, Wölfel R. 2022. Clinical and virological features of first human monkeypox cases in Germany. Infection https://doi.org/10.1007/s15010-022-01874-z.

20. Erez N, Achdout H, Milrot E, Schwartz Y, Wiener-Well Y, Paran N, Politi B, Tamir H, Israely T, Weiss S, Beth-Din A, Shifman O, Israeli O, Yitzhaki S, Shapira SC, Melamed S, Schwartz E. 2019. Diagnosis of Imported Monkeypox, Israel, 2018. Emerg Infect Dis 25:980–983.

21. Platten M, Hoffmann D, Grosser R, Wisplinghoff F, Wisplinghoff H, Wiesmüller G, Schildgen O, Schildgen V. 2021. SARS-CoV-2, CT-Values, and Infectivity-Conclusions to Be Drawn from Side Observations. Viruses 13:1459.

22. 2022. Lab Alert: MPXV TNF Receptor Gene Deletion May Lead to False Negative Results with Some MPXV Specific LDTs. https://www.cdc.gov/locs/2022/09-02-2022-lab-alert-MPXV_TNF_Receptor_Gene_Deletion_May_Lead_False_Negative_Results_Some_MPXV_Specific_LDTs.html. Retrieved 2 September 2022.

23. 2022. Select Agents and Toxins List | Federal Select Agent Program. https://www.selectagents.gov/sat/list.htm. Retrieved 5 September 2022.

